# AXIS: A Lab-in-the-Loop Machine Learning approach for generalized detection of macromolecular crystals

**DOI:** 10.1101/2025.11.03.685844

**Authors:** Aurelien Personnaz, Sihyun Sung, Raphael Bourgeas, Sruthi Sunni, Florine Dupeux, Bukunmi Adediran, Rosicler Barbosa, Anne-Sophie Humm, Euan Colaco-Osorio, José Antonio Márquez

**Affiliations:** European Molecular Biology Laboratory, 71 Avenue des Martyrs, Grenoble, 38000, France; Institut de Biologie Structurale, CNRS, 71 Avenue Des Martyres, Grenoble, 38000, France

**Keywords:** Structural Biology, AI, Lab in the Loop, Image processing, Automation, high throughput crystallography

## Abstract

Macromolecular crystallography provides mechanistic understanding of biological processes and can be applied in drug design. Nowadays, the use of robotic systems for crystal growth and diffraction analysis is widespread and high throughput protein-to-structure pipelines for ligand and fragment screening are revolutionizing the field. However, the identification of crystals is still largely carried out through manual inspection, sometimes involving tens of thousands of images, which represents a bottleneck in an otherwise highly automated process. Here we describe **AXIS**, an **A**I-based **C**rystal **I**dentification **S**ystem combining the DINOv2 computer vision model, state-of-the-art transfer learning and MARCO, the largest crystallization dataset available to date, for automated crystal detection. AXIS can operate both with visible and UV light images and integrates a Lab-In-The-Loop approach combining ML and expert inputs for continuous learning and specialization. AXIS enables automated annotation of large crystallization image datasets with performance and accuracy comparable to that of human experts and the Lab-In-The-Loop approach introduced here enables efficient adaptation to local conditions facilitating widespread application, which has been a major limitation to date. AXIS can help correct human errors in image annotation and removes critical bottlenecks, particularly in the context of extensive crystallization screens or high throughput applications like fragment and ligand screening unlocking the potential for higher levels of automation that are key both in fundamental and translational research.

**Graphical Abstract:** 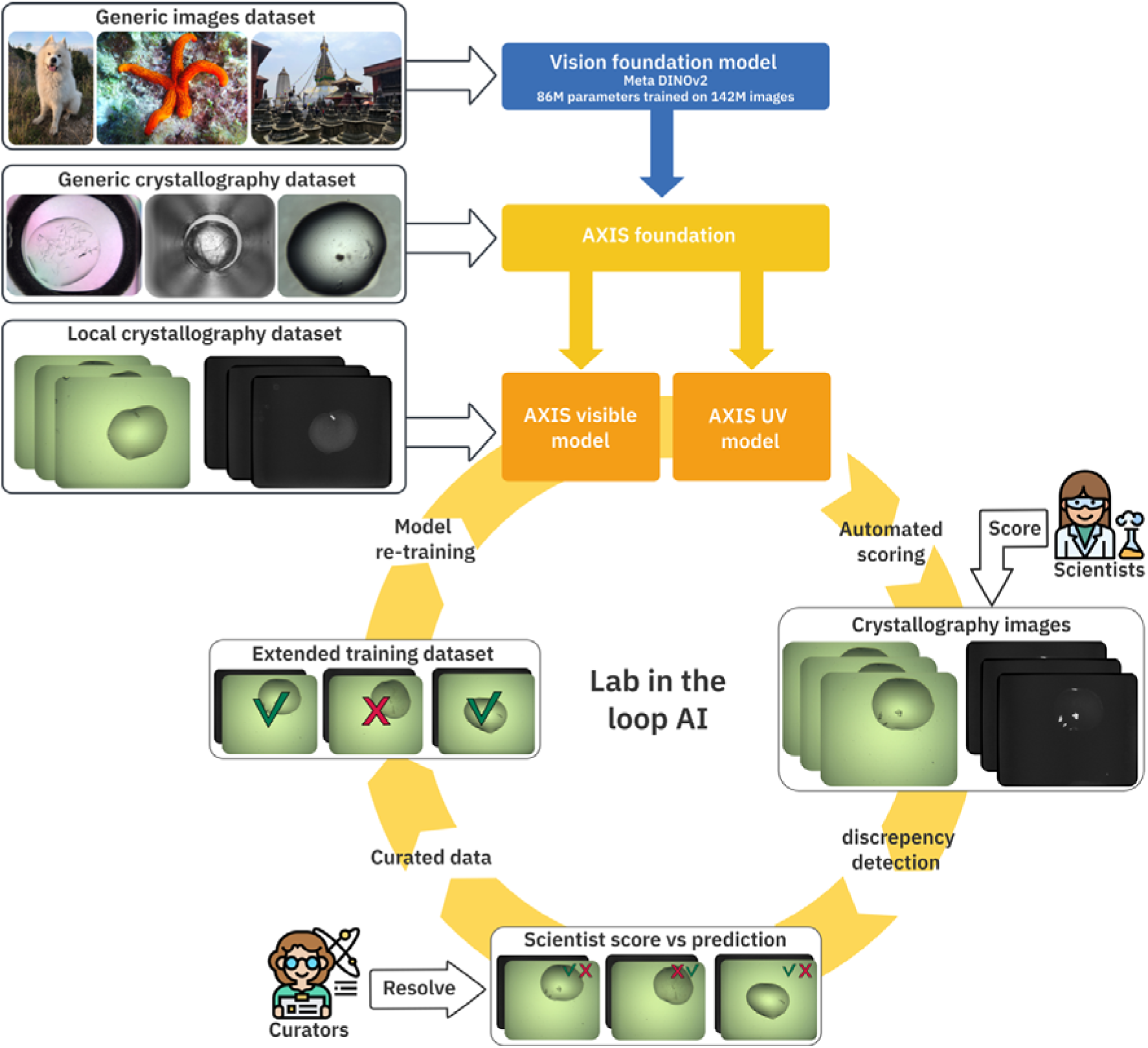

## 1. Introduction

Macromolecular Crystallography (MX) along with other techniques like CryoEM, NMR or AI-based fold predictions can be applied to the study of protein structure and function providing mechanistic understanding of biological processes. This contributes fundamental knowledge that underpins our understanding of health and disease states and can inform the development of novel therapies and applications in biotechnology (Helliwell, 2017; Whittle & Blundell, 1994). Automation has been introduced at nearly all steps of the MX experimental workflow, from crystallization to diffraction data collection and processing and highly automated protein-to-structure pipelines are currently available (Rupp *et al*., 2002; Cipriani *et al*., 2006; Cusack *et al*., 1998; McCarthy *et al*., 2018; Bowler *et al*., 2015; Zander *et al*., 2016; Cornaciu *et al*., 2021; Schwalbe *et al*., 2024). Recently, new technology developments have enabled high throughput X-ray based ligand and fragment screening democratizing access to structure-based drug design approaches (Cox *et al*., 2016; Thomas *et al*., 2019; Cornaciu *et al*., 2021; Münzker *et al*., 2020; Barthel *et al*., 2024), as well as the study of dynamic and time-resolved systems (Schlichting, 2015; Stauch & Cherezov, 2018; Mehrabi *et al*., 2019).

In any of its applications, MX requires generation of crystals with high diffracting power. This is typically a complex process that involves performing and evaluating large numbers of crystallization experiments. The most commonly used crystallization technique involves the so-called *vapour diffusion* method, in which a supersaturated solution of the biological molecule is achieved by adding mild precipitant agents under different chemical and physical condition, eventually leading to the formation of crystals (McPherson, 2017). It is difficult to predict crystallization conditions for molecules that have never been crystallized before, therefore screening of different types precipitants, like salts and polymers against different experimental parameters like pH, ionic strengths, temperatures etc, is necessary leading to a large number of crystallization experiments that need to be evaluated individually (McPherson, 2017; Newman *et al*., 2005). Moreover, crystals may take from hours to days or even weeks to develop under vapour diffusion conditions. During this period each experiment has to be visually inspected multiple times for the appearance of crystalline material. The initial screening experiments often help identify starting crystallization conditions that need to be optimized through an iterative process involving new crystallization experiments and visual inspections. On the other hand, applications like ligand and fragment screening require the identification of hundreds of crystals. Therefore, crystallography projects require regular inspection of hundreds to thousands of crystallization experiments.

Nowadays, crystallization is typically carried out with the use of specialized robots and in 96-well microplate format (Newman *et al*., 2005). Automated crystal farms that hold from dozens to a thousand 96-well crystallization plates at controlled temperatures, take images of each crystallization drop at regular intervals (Chayen & Saridakis, 2008; Stock *et al*., 2005; Dimasi *et al*., 2007; Cornaciu *et al*., 2021). As an example, at the Hight Throughput Crystallization Facility at EMBL Grenoble, a typical crystallization screening experiment involves 6 96-well plates in which 576 different crystallization conditions are tested for a sample at three different concentrations. This results in 1.728 individual experiments that are automatically imaged in our crystal farms 9 times during a period of several weeks (Dimasi *et al*., 2007; Cornaciu *et al*., 2021). Similarly, fragment screening projects involve generation of hundreds to thousands of crystallization experiments (Münzker *et al*., 2020; Cornaciu *et al*., 2021). Images, along with crystallization parameters are harvested by the Crystallographic Information Management System (CRIMS) (Cornaciu *et al*., 2021; Healey *et al*., 2021) and presented to the user through a web-based interface for evaluation of results. Therefore, a typical experiment will result in 15.552 images that have to be individually evaluated. Due to the iterative nature of this approach, scientist will carry out multiple such experiments in the course of a project varying parameters, like temperature, construct design etc either successively or in parallel, which may result in hundreds of thousands of images. Similar numbers of experiments are produced at other crystallization facilities around the world. A non-exhaustive catalogue of crystallization facilities can be found at https://instruct-eric.org/platform-type/x-ray-techniques.

Currently, identifying crystals in crystallization images relies largely on visual inspection and can be a difficult task. Crystals can show very different morphologies and may appear along with amorphous precipitate, phase separation or other type of objects partially masking them. On the other hand, very small or microcrystalline material may be difficult to detect with images taken from automated systems. Therefore, even well trained scientists may fail to recognize crystals either through inherent difficulties or through a decrease in attention over extended evaluation periods (Bruno *et al*., 2018). Indeed, it has been shown that when presented with the same set of crystallization images, multiple crystallographers may disagree in their scores as much as 30 % of the images (Bruno *et al*., 2018; Wilson, 2002). A number of computational tools for automated scoring of crystallization micrographs have been proposed using various approaches and with different levels of performance (Wilson, 2002; Cumbaa & Jurisica, 2010; Cumbaa *et al*., 2003; Bern *et al*., 2004; Saitoh *et al*., 2005; Liu *et al*., 2008; Pan *et al*., 2006). In a landmark collaborative study, the Machine Recognition of Crystallization Outcomes (MARCO) initiative, a large dataset containing 493,214 crystallization images from 5 different laboratories was assembled. (Bruno *et al*., 2018; Rosa *et al*., 2023). These images were scored in 4 classes (crystal, precipitate, clear and other) and a deep Convolutional Neural Network (CNN) using the Inception-v3 architecture was trained for automated classification achieving 91% crystal recall and 94% accuracy over all the classes (Bruno *et al*., 2018). More recently, a new crystal image classifier, CHiMP-V2 was developed (King *et al*., 2024). In this work, the ConvNeXt-Tiny image vision model (Liu *et al*., 2022) was trained first on the MARCO dataset and then on a smaller set of 11.167 local images from the Diamond Light Source VMXi beam line (Sanchez-Weatherby *et al*., 2019b). This classifier used the same categories as the MARCO model and achieved crystal recall of 82% to 90 % with precision of 65% to 70% (King *et al*., 2024). In a modified version of this system, the CHiMP detector, the four categories were collapsed into two, crystals and no-crystals achieving between 92% and 95% crystal detection rates but with lower precision, 33% to 44 % (King *et al*., 2024). However, it has been found that performance of these models decreases when applied to local datasets. At the same time, performance can improve by retraining the model with a limited number of images generated locally (Rosa *et al*., 2023). However, many facilities lack the resources and expertise to collect, annotate and train ML models to adapt to local conditions. Interestingly none of these systems explored the use of UV imaging for Machine Learning (ML) based crystal detection, although UV-capable imaging systems have become common in many crystallization facilities.

In ML classification, recall and precision (see materials methods section for definitions) are correlated and require balancing for optimal performance. Crystal identification requires very high recall (low number of false negatives). Recall can typically be increased at the expense of precision, but the lower the precision, the larger the number of false positives introduced, decreasing the value of the classifier. This is particularly important as crystallization experiments are highly imbalanced, with a very high proportion of images containing no crystals (95% or more) potentially leading to high number of false positives when precision is moderate. Two additional problems of crystallization image classification tools have been the requirement for large training datasets and the loss of performance when applied to images originating from local infrastructures. Crystallography facilities use different types of crystallization plates, imaging equipment and apply different illumination and imaging settings for example, all of which, can affect performance in ML classification (Rosa *et al*., 2023).

Recent developments in AI have revolutionized the field of computer vision including the development of vision transformers (Dosovitskiy *et al*., 2020) large computer vision models (Oquab *et al*., 2023) and advanced transfer learning techniques (Hu *et al*., 2021a; Zhuang *et al*., 2020) opening new opportunities for crystallization image classification. In this work we present AXIS, AI-based Crystal Identification System, integrating modern computer vision models, state-of-the-art transfer learning techniques and a Lab-In-the-Loop approach for continuous learning. This system can evaluate both visible and UV crystallization images and achieves very high performance across datasets from different origins. The CRIMS web-based Lab-In-The-Loop module makes it possible to integrate federated input from expert crystallographers for continuous fine-tuning based on new experimental data. This approach provides a foundational model for crystal identification and enables rapid adaptation to local conditions with minimal effort, helping facilities obtain best performance under their specific conditions.

## 2. Material & Methods

### 2.1. Training and test datasets

Different training and test datasets were either used or generated in this work and are presented in Table 1. Images from the training and test sets were always kept separated and those from the test sets were never use for training. For our first step in the transfer learning process (see below) we used a training set combining the MARCO and C3-Suplementary training datasets. The MARCO dataset was generated and published by the Machine Recognition of Crystallization Outcomes (MARCO) initiative (Bruno *et al*., 2018). Five crystallography facilities (Collaborative Crystallisation Centre, GlaxoSmithKline, Hauptman-Woodward Medical Research Institute, Merck & Co., Bristol-Myers Squibb) collaborated to gather 462 804 crystallography outcome micrographs in the visible spectrum with labels from four different classes (crystal, clear, precipitate, other). Here we only used the images from the MARCO training set (415 777 visible images). The Collaborative Crystallisation Centre (C3) extended this dataset with an additional training set of 30 767 images (Rosa *et al*., 2023; Rosa & Newman, 2021) distributed in the same classes. Here we combined these two datasets into a single training set and consolidated it into two classes: “crystals”, containing the images labelled as crystals and “other” containing images labelled as clear, precipitate or other in the MARCO and C3-Suplemnatery datasets. For simplicity we will call this new extended dataset the MARCO-C3 dataset and it is composed of 76 836 and 369 708 in the “crystal” and “other” classes respectively See table 1.

**Table 1.** Training and test datasets used and generated in this work.

| Dataset name | Training/Test | Images (Visible)<br>Crystal / Other | Images (UV)<br>Crystal /Other | Reference | DOI |
| --- | --- | --- | --- | --- | --- |
| MARCO-C3 | Training | 76 836 / 369 708 |  | Bruno et al.(Bruno *et al.*, 2018) | https://ubir.buffalo.edu/xmlui/handle/10477/77793 |
| C3-Test | Test | 2 509 / 6 359 | 2 509 / 6 359 | Rosa et al.(Rosa *et al.*, 2023) | https://doi.org/10.5281/zenodo.4635300 |
| CRIMS-test | Test | 225 / 3 111 | 225 / 3 111 | This work | https://doi.org/10.5281/zenodo.17279081 |
| CRIMS-v1 | Training | 4 707 / 3 171 | 4 707 / 3 171 | This work | https://doi.org/10.5281/zenodo.17279591 |
| CRIMS-v2 | Training | 5 934 / 19 634 | 5 293 / 11 365 | This work | https://doi.org/10.5281/zenodo.17279968 |
| CRIMS-v3 | Training | 13244 / 105 094 | 6153 / 43499 | This work | https://doi.org/10.5281/zenodo.17426047 |
| VMXi-training | Training | 6 752 / 4 409 |  | King et al.(King *et al.*, 2024) | https://doi.org/10.5281/zenodo.11097395 |
| VMXi-test | Test | 145 / 487 |  | King et al.(King *et al.*, 2024) | https://doi.org/10.5281/zenodo.11097395 |

Along with the C3-Suplemnatery training set, the Collaborative Crystallisation Centre published the C3-test set (Rosa *et al*., 2023; Rosa & Newman, 2021) that we also used here for performance evaluation. This set is composed of both visible and UV light micrographs, although UV images were not exploited for automated classification in the original work. The C3-test set was also consolidated into two classes as described above with 2 509 images in the “crystals” and 6 359 “other” classes respectively. We kept the original name for this set, C3-test (Table 1).

We generated a test set, CRIMS-test, with local images from the HTX facility in EMBL Grenoble (Cornaciu *et al*., 2021) to evaluate performance of the different classifier models with local images (Table 1). These images were generated with two Rock Imager 1000 imaging robots (Formulatrix, Bedford, MA, United States) equipped with visible and UV imaging systems. Crystallization experiments were set up in 96-well format using CrystalDirect plates (SKU: M-XDIR-96-3-40, Mitegen, Ithaca, NY USA) as described previously (Dimasi *et al*., 2007; Cornaciu *et al*., 2021; Healey *et al*., 2021; Zander *et al*., 2016). Particular care was taken to ensure maximal diversity and avoid potential redundancy in this set. For this, random sampling of images from crystallization plates in the historical CRIMS experimental database was carried out. Only crystallization plates corresponding to initial crystallization screening experiments were considered. Crystal optimisation plates were excluded from this selection to prevent overrepresentation of certain crystal types. Only one imaging session was selected from each of the plates to eliminate the possibility of including images from the same experiment taken at different time points, which might be redundant otherwise. The imaging session used was chosen to correspond to a dual Visible/UV imaging to facilitate evaluation of both visible and UV based classifiers with the same test set. The selected plates represent a random sampling from different projects brought by our users and processed at the facility over a period of 3 years. Images were manually annotated by multiple expert operators and when agreement between scorers could not be reached, they were discarded. This resulted in the CRIMS-test set with 3 335 pairs of visible and UV images including 231 and 3 104 images in the crystal” and “other” classes respectively (Table 1).

Three additional training datasets, CRIMS-v1, CRIMS-v2 and CRIMS-v3 were generated with images extracted from the CRIMS experimental database to support fine tuning of the CRIMS-AXIS classifiers and the Lab-In-The-Loop training process. The CRIMS-v1 training dataset consists of 7 878 pairs of visible and UV light micrographs, composed of 4 707 “crystals” and 3 171 “others”. In order to increase the representation of crystals in the initial local training set, the CRIMS database was queried for images with annotations corresponding to the crystals and these annotations were then validated by experts.

The CRIMS-v2 training dataset was generated as part of the Lab-In-The-Loop process (see below). The CRIMS-AXIS-v1 classifier (Table 2) was applied to automatically annotate crystallization experiments generated at the EMBL HTX Lab in Grenoble. A series of crystallization plates were randomly selected from the CRIMS experimental database as described for the CRIMS Test set above. Results of ML (AXIS) and manual scoring were compared for those plates. Experiments showing discrepancies between the human and CRIMS-AXIS-v1 scoring were subjected to a curation process in which experts individually evaluated the images and decided on the correct annotation for the experiment that were then included in the CRIMS V2 dataset. Images that could not be unambiguously assigned by the experts to one or other class were discarded. Experiments showing agreement between the human and AXIS scoring were also included in the dataset. In total the CRIMS V2 training dataset contains 25 568 pairs of visible and UV light images including 5 934 and 19 634 images in the “crystal” and “other” classes respectively. The CRIMS-v3 training dataset was generated through a new iteration of the Lab-In-The-Loop process, but in this case using the AXIS-CRIMS-v2 model for automated annotation (Table 1).

**Table 2.** Performance of AXIS crystal identification models.

| Model | CRIMS-test set |  | C3-test set |  |
| --- | --- | --- | --- | --- |
|  | Balanced accuracy | Recall | Balanced accuracy | Recall |
| MARCO | 83.00% | 67.41% | 90.70% | 88.40% |
| AXIS-foundation | 89.37% | 81.25% | 90.86% | 98.80% |
| AXIS-CRIMS-v1 | 91.87% | 95.54% | 90.20% | 98.68% |
| AXIS-CRIMS-v2 | 95.34% | 95.98% | 91.16% | 98.53% |
| AXIS-CRIMS-v3 | 96.27% | 96.43% | 93.11% | 97.69% |

Finally, to assess the reproducibility of our system with images from other sources, we used training and test datasets published by the Diamond light source(King *et al*., 2024), including the VMXi beam line classification training dataset composed of 6 752 “crystal” and 4 409 “other” images, and the VMXi unambiguous test set, composed of 145 “crystal” and 487 “other” images. For both of these datasets, the images were taken with visible light imaging.

### 2.2. Machine learning

Previous works on AI applied to crystallization micrograph classification employed the Convolutional Neural Network (CNN) architecture (Bruno *et al*., 2018; King *et al*., 2024), which used to be the foundation of computer vision for many years. In the late 2010s, a major breakthrough happened in AI with the development of transformer networks(Vaswani *et al*., 2017). At first designed for Natural Language Processing, they leveraged the “attention” concept to understand and learn languages underlying logic and allowing the emergence of modern Large Language Models (Devlin *et al*., 2018) (LLMs). These concepts were also rapidly adapted to the field of computer Vision with the appearance of vision transformers (Dosovitskiy *et al*., 2020). While more demanding computationally, those large models proved more efficient than CNNs when trained on sufficiently large datasets (Dosovitskiy *et al*., 2020). They also were also shown to generalize better than CNNs to downstream tasks with fine-tuning (Zhou *et al*., 2021). After initial evaluation of several of the computer vision models recently published we selected the DINOv2-base model that we obtained from Hugging Face repository (https://huggingface.co/facebook/dinov2-base). The DINOv2(Oquab *et al*., 2023) model was developed by Meta AI as a multi-purpose foundation model for Vision tasks. Applying similar self-supervision techniques used to train LLMs on vast amounts of non-labeled data (Devlin *et al*., 2018), DINOv2 was trained in a self-supervised way on 142 million of non-labelled images, allowing it to obtain a task-agnostic understanding of visual features and image analysis(Oquab *et al*., 2023).

The idea behind transfer learning or fine-tuning (Zhuang *et al*., 2020) is to benefit from the projection capabilities a model has learned while solving an initial problem, generally with large amounts of data available, and adapt it to a comparable or more specific task for which limited data is available. Fine-tuning the entirety of a very large network like DINOv2, with 86 million parameters, would require a very high volumes of data and computational power to be efficient. The traditional way to fine-tune such models consists in re-training only the classification head (upper levels of the neural network). This way the mathematical projection part of the model is not modified, and only the classification of the projections is trained. This method simplifies greatly the training, but limits the level of adaptation to the new data. An alternative recently proposed is Low Rank Adaptation (Hu *et al*., 2021a) (LoRA). With LoRA, low-rank weight matrices are injected to certain parts of the model (typically attention and feedforward layers), while the larger part of the original model remains untouched. During training, only these low-rank matrices are updated. This technique limits the number of parameters to be trained, while at the same time allows to apply the fine-tuning process across all layers of the model. After the training, the low rank matrices can either be merged into their target layers to make their changes permanent, or they can be kept as LoRA adapters to be further trained. For this work we used the LoRA implementation from Hugging Face Parameter Efficient Fine Tuning (https://github.com/huggingface/peft). We injected matrices into all dense layers from the model, with an alpha parameter value of 20 and a rank of 25 (respectively the scaling factor of the matrices on the original weights, and the rank of the low-rank decomposition), resulting in the training of four million parameters instead of the original eighty-six million (Table 7). We kept the matrices in separated adapters, allowing us to run two successive fine tunings on them.

The first fine-tuning was from the original DINOv2-base on the MARCO-C3 dataset composed of 446 344 images. It was applied using a single Nvidia A100 GPU on the EMBL HPC cluster in Heidelberg. We used Hugging Face trainer API (a python framework based on torch). We ran two epochs with a linearly decreasing learning rate starting at 0.0005 and batches of 32 images. To handle the systematic imbalance of crystallography datasets, weighted cross entropy loss was used with inverse frequency weighting to define the classes weights. The images were normalized and resized to 518x518 pixels as the original DINOv2 training set. They were then turned to grayscale, and minimal augmentations were applied (vertical and horizontal flipping) as crops could possibly cut crystals from the images, and shape transformations could alter their characteristic geometric features. This first training took 12 hours, and resulted in the AXIS-foundation model (see Tables 1 and 2). Additional fine-tuning steps were carried out to increase performance with images from our local infrastructure as described in the results section. These were executed in the same way as indicated above, although a lower end Nvidia RTX 2080 TI was used with a batch size lowered to 16 to fit into the smaller memory, and the trainings were done with only one epoch. These fine tunings were independently run on visible light and UV light images, resulting in separate models (see Tables 1 and 2). Each of those fine tunings took 45 minutes to one hour of computing time. For comparison purposes, the previously established MARCO (Bruno *et al*., 2018)and Chimpv2(King *et al*., 2024) models were obtained from https://github.com/tensorflow/models/tree/master/research/marco and https://doi.org/10.5281/zenodo.11190973 respectively.

### 2.3. Metrics

The evaluation of performance of automated scoring models for crystallization outcomes requires metrics that are appropriate for highly imbalanced datasets. The main metrics used in this work were crystal recall, called here simply “recall”, defined as the proportion of crystal events correctly labelled, and balanced accuracy. Both are critical as high crystal recall with low accuracy would be detrimental due to the overabundance of images in the “other” class. The most commonly reported metric in classification tasks is accuracy, defined as the proportion of correctly classified instances. Accuracy is straightforward to calculate and widely used (Bruno *et al*., 2018; Rosa & Newman, 2021). However, it can be misleading in highly imbalanced datasets—such as those in crystallography, where actual crystals are rare. For example, a model that predicts “no crystal” for every drop would achieve high accuracy simply because the majority class, “no crystals”, is much larger, yet it would fail to detect the very events of interest. The F1 score, which combines precision (the fraction of predicted positives that are truly positive) and recall (the fraction of true positives that are detected) into a single harmonic mean has also been used (King *et al*., 2024). While the F1 score is more sensitive to minority classes than accuracy, it still depends heavily on the underlying distribution of classes in the test dataset. If the proportion of crystal images varies across datasets, direct comparison of F1 scores can become problematic.

In this context, balanced accuracy provides a more stable measure. Balanced accuracy averages the recall scores across classes, giving equal weight to the minority and majority classes. This ensures that even if crystals represent only a small fraction of all drops, their detection is as influential to the metric as the much larger class of non-crystal outcomes. Balanced accuracy is therefore better suited than both raw accuracy and the F1 score when the goal is to identify rare but meaningful events.

## 3. Results and Discussion

### 3.1. Applying the DINOv2 computer vision model and transfer learning for automated classification of crystallisation images

The field of AI is evolving very rapidly and revolutionizing research in biology(Abramson *et al*., 2024; Jumper *et al*., 2021; Yu *et al*., 2023; Krishna *et al*., 2024; Baek *et al*., 2021). Large language models have transformed text analysis and a similar revolution has taken place in computer vision. Just as language models learn grammatical rules and vocabulary from massive text (Vaswani *et al*., 2017; Devlin *et al*., 2018), modern “foundation models” in computer vision are trained on millions of generic images, learning to recognize shapes, textures, and structures to provide an accurate numerical description of the content of the image (Dosovitskiy *et al*., 2020; Oquab *et al*., 2023). Once trained on large numbers of generic images, these models can be re-trained, or fine-tuned, to perform a more specific task with relatively little additional data through a process known as transfer learning (Zhuang *et al*., 2020; Devlin *et al*., 2018). We wanted to investigate whether the DINOv2 computer vision model, trained on 142 million curated natural images and recently released (Oquab *et al*., 2023), could be applied to automatically identify crystals within micrographs in the context of high throughput macromolecular crystallization experiments. At the same time, we wanted to test whether UV imaging could contribute to crystal identification. UV imaging has become common in high throughput crystallization laboratories, but has not yet been systematically applied in the context of ML-based crystal identification.

To make the best use of modern foundation models and publicly available data while minimizing the costs and complexity, we designed a multi-step training process based on successive transfer learning steps to obtain our classification models (Figure 1). The process starts with the DINOv2 foundation model pre-trained on generic and diverse images to identify and understand visual features (Oquab *et al*., 2023). We then combined the previously published MARCO and C3 image datasets (Bruno *et al*., 2018; Rosa *et al*., 2023) to re-train this model for the identification of crystals within crystallization images obtained with visible light (See materials and methods section). The MARCO-C3 training dataset contains 446 544 images distributed across 4 class (Clear, Crystal, Precipitate and other). We consolidated this dataset into two classes, *crystals* containing 76.936 images and *other* with 369.708 images and then re-trained the DINOv2 model for two-class image classification. Similarly, the C3 test dataset(Rosa *et al*., 2023) was consolidated into two classes (2.509 *crystals*, 6.359 *other*) and was used to evaluate the performance of the trained models (See material and methods section and Table 1).

**Figure 1.**
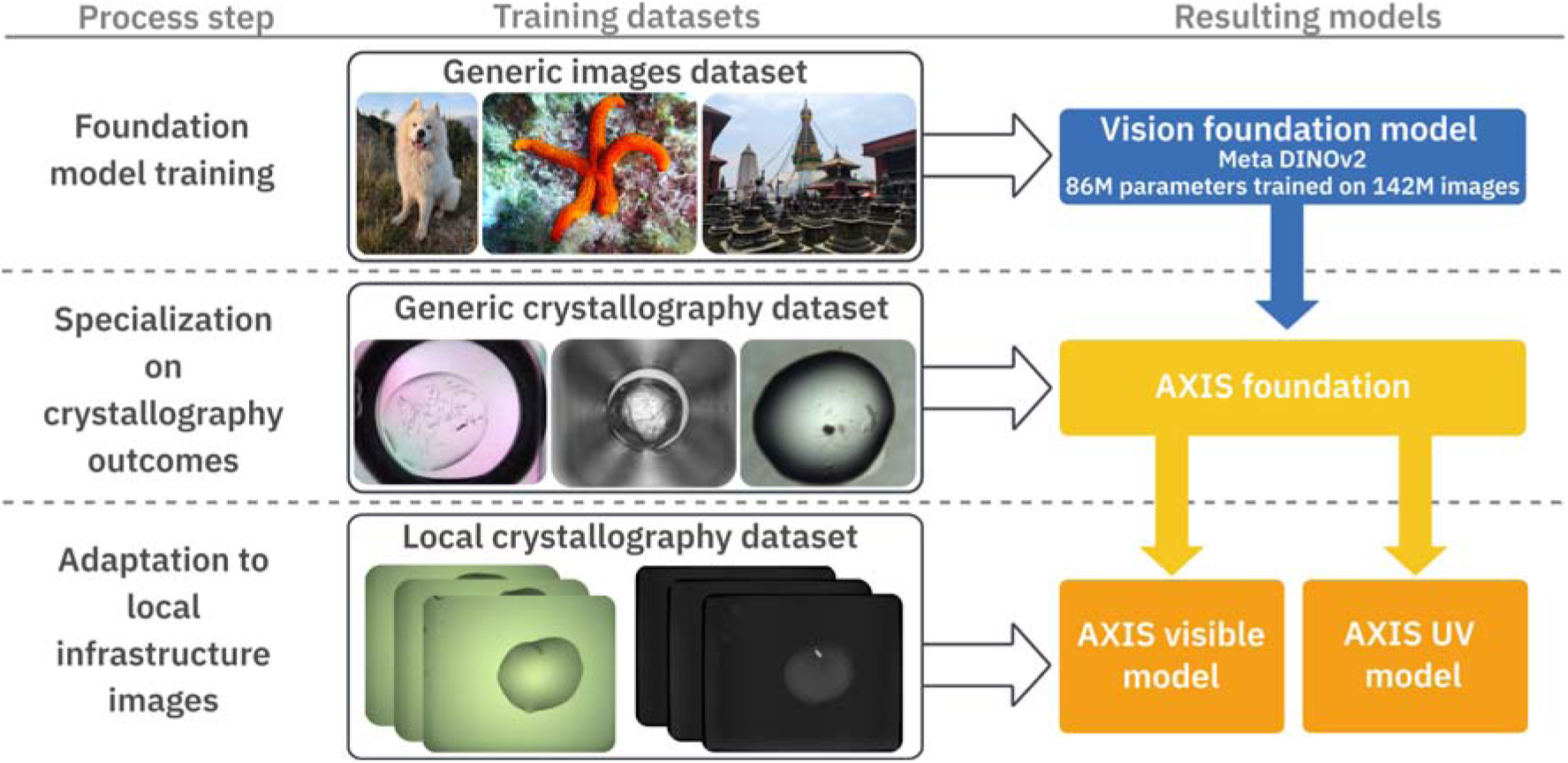
Schematic representation of the AXIS multi-step transfer learning process. The DINOv2 model trained on 142 M images was used as starting point (top). LoRA transfer learning on this model was applied using visible light images from the MARCO training dataset to generate the AXIS Foundation model (middle). This model was then independently re-trained with Visible and UV images to generate two independent classifiers that operate in Visible and UV images respectively (bottom panel).

Transfer learning with very large AI models can be challenging and may require significant computational resources. For example, the DINOv2 model is a large transformer network composed of 86 million parameters distributed in a repetition of encoder blocks with multi-head self-attention and feed-forward fully connected sub-networks. The traditional transfer learning method consisted in replacing the classification head while blocking the training of every other layer of the foundational model (Yosinski *et al*., 2014). This allowed to preserve the projection capability of the foundation model (obtain the most meaningful mathematical representation from any input), while quickly re-training the classification part – i.e. the model task. However, while allowing re-training in an acceptable time frame with limited amounts of data, this technique prevented any adaptation of the projection part– or encoding –of the original model, limiting the scale of the re-training. An alternative was therefore introduced with LoRA: Low-Rank Adaptation of Large Language Models(Hu *et al*., 2021b). The idea behind LoRA is also to freeze the foundational model weights, but to inject low-rank weight matrices to certain layers of the model (typically attention or feedforward layers). During training, only these low-rank matrices are updated, allowing to train only a limited number of parameters with improved efficiency, while extending the adaptation to the whole model to efficiently specialize large models on a given task.

By re-training the DINOv2 model with the MARCO-C3 image dataset (see materials and methods) we obtained the first AXIS crystal identification model, AXIS-Foundation. To evaluate the performance of the training process we used two different test image datasets. The C3 test dataset (Rosa *et al*., 2023) (with 2.509 and 6.359 images in the Crystal and other class respectively) and the CRIMS test set, with 3.335 images ( 231 crystals, 3104 other) produced at the EMBL High Throughput Crystallization facility in Grenoble (Dimasi *et al*., 2007; Cornaciu *et al*., 2021) (Table 1). For the later, images were randomly selected from the CRIMS database corresponding to different types of proteins and complexes and representative of a wide variety of structural biology projects. Care was taken to eliminate redundancy, for example excluding multiple imaging sessions from the same experiment (see materials and methods section) and only primary screening experiments were included. The distribution of images among the two classes in the CRIMS test dataset also reflects that of typical crystallization screening projects. We evaluated the performance of both AXIS-Foundation and the previously published MARCO model against these two test datasets. As can be observe in Table 2, AXIS Foundation produced better results as compared with MARCO for both the C3 and CRIMS test sets. For the C3 test dataset, the AXIS-Foundation model showed higher level of crystal recall, 98.8 % versus 88.4% for the original MARCO model, with similar levels of accuracy (Table 2). However, when applied to the CRIMS Test dataset both MARCO and AXIS Foundation showed decreased performance. This confirmed the previous observation that the performance in crystal image classification tends to decrease when applied to local images, (Rosa *et al*., 2023). This might be due to a number of factors, like differences in plate types, imaging equipment, or imaging and illumination settings for example, leading to differences in global visual features. We decided to test whether re-training the AXIS-Foundation model with local images (obtained at our facility) would improve performance, but at the same we wanted to explored whether the introduction of UV images could help in improving crystal identification.

### 3.2. Combining visible and UV imaging for crystal detection

In the later years commercial systems for automated crystal imaging under both visible and UV light have been introduced and are currently in use in many laboratories. Usually, these systems exploit the intrinsic fluorescence of proteins. We set out to confirm whether the addition of local visible images in the training process would improve the performance of the AXIS system but also whether the use of UV images could help in improving automated crystal detection. For this purpose, we generated an in-house training dataset with experiments extracted from the CRIMS database, the CRIMS-v1 training set (Table 1). This training dataset contains 7.878 pairs of visible and UV images (4.707 crystals, 3.171 other) selected from a diversity of historic user projects supported at the HTX facility in Grenoble. We selected this dataset so that for every visible image there would also be an equivalent image taken with UV light during the same imaging session only a few minutes apart. Similarly, and to facilitate performance evaluation, the CRIMS test set, described above that was never used for training, was selected to contain both visible and UV images from the same imaging session (see materials and methods). The LoRA approach was applied again to retrain the AXIS-Foundation model with the CRIMS-v1 training dataset. We decided to generate two completely independent classification models for visible and UV images respectively, as both types of images are not always available for the same experiment, which means that in many cases only one or the other classifier would need to be applied. As a result, two independent classification models were generated, AXIS-Vis-v1 and AXIS-UV-v1. As can be observed in Table 3, the AXIS-Vis-v1 showed improved performance as compared to the AXIS-Foundation model, achieving crystal recall of 90.0% and balanced accuracy of 93.1% on the CRIMS test set. This confirms that addition of a limited number of local images in the training set improves performance in crystal classification.

**Table 3.**
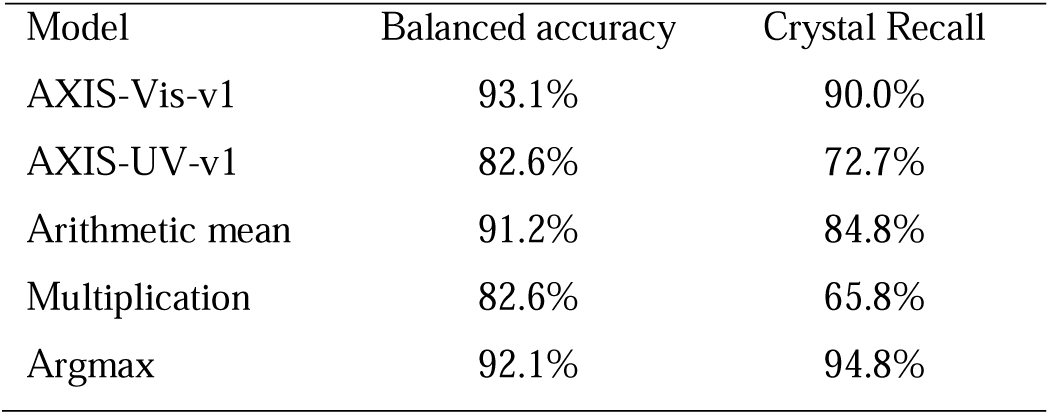
Performance of visual and UV-based image classifiers and several output aggregation methods on the CRIMS-test dataset.

**Table 4.** Visual and UV light images prediction aggregation method comparison for AXIS-CRIMS-v1 on CRIMS-test.

| Model | Accuracy | Balanced accuracy | Recall | Precision |
| --- | --- | --- | --- | --- |
| Visual only | 95.6% | **93.1%** | 90.0% | 63.0% |
| UV only | 91.1% | 82.6% | 72.7% | 41.7% |
| Arithmetic mean | 96.8% | 91.2% | 84.8% | 83.0% |
| Multiplication | **97.1%** | 82.6% | 65.8% | **89.9%** |
| Argmax | 89.7% | 92.1% | **94.8%** | 39.7% |

The AXIS-UV-v1 model, on the other hand, achieved moderate performance, with a crystal recall of 72.7% and a balanced accuracy of 82.6%. This is not unexpected, as the MARCO dataset used here did not contain UV data and hence this model has been trained with a considerably lower number of images. Notably, the AXIS-UV-v1 model produced relatively high number of false positives in the crystal class. This might be in part due to the fact that often, amorphous protein precipitates show strong fluorescence signals that could be confused with signal from crystals. Interestingly, despite its moderate performance, the AXIS-UV-v1 classifier was able to identify crystals that were not recalled by the AXIS-Vis-v1 model. This is exemplified in Figure 2. Figure 2a shows a visible image with crystals that are obscured by a layer of protein precipitate and that were missed by the visible image classifier. However, the corresponding UV image (Figure 2b) shows clear UV signals for these crystals and the UV-based classifier correctly assigned this image to the *crystal* class. Similarly, panels e-f in figure 2, show very small microcrystals with poor contrast under visible light but that were readily recognised by the UV model. Conversely, Panels c-d and g-h of Figure 2 show crystalline material identified by the visible classifier but that either showed insufficient signal or resolution under UV light and were missed by the AXIS UV model.

**Figure 2.**
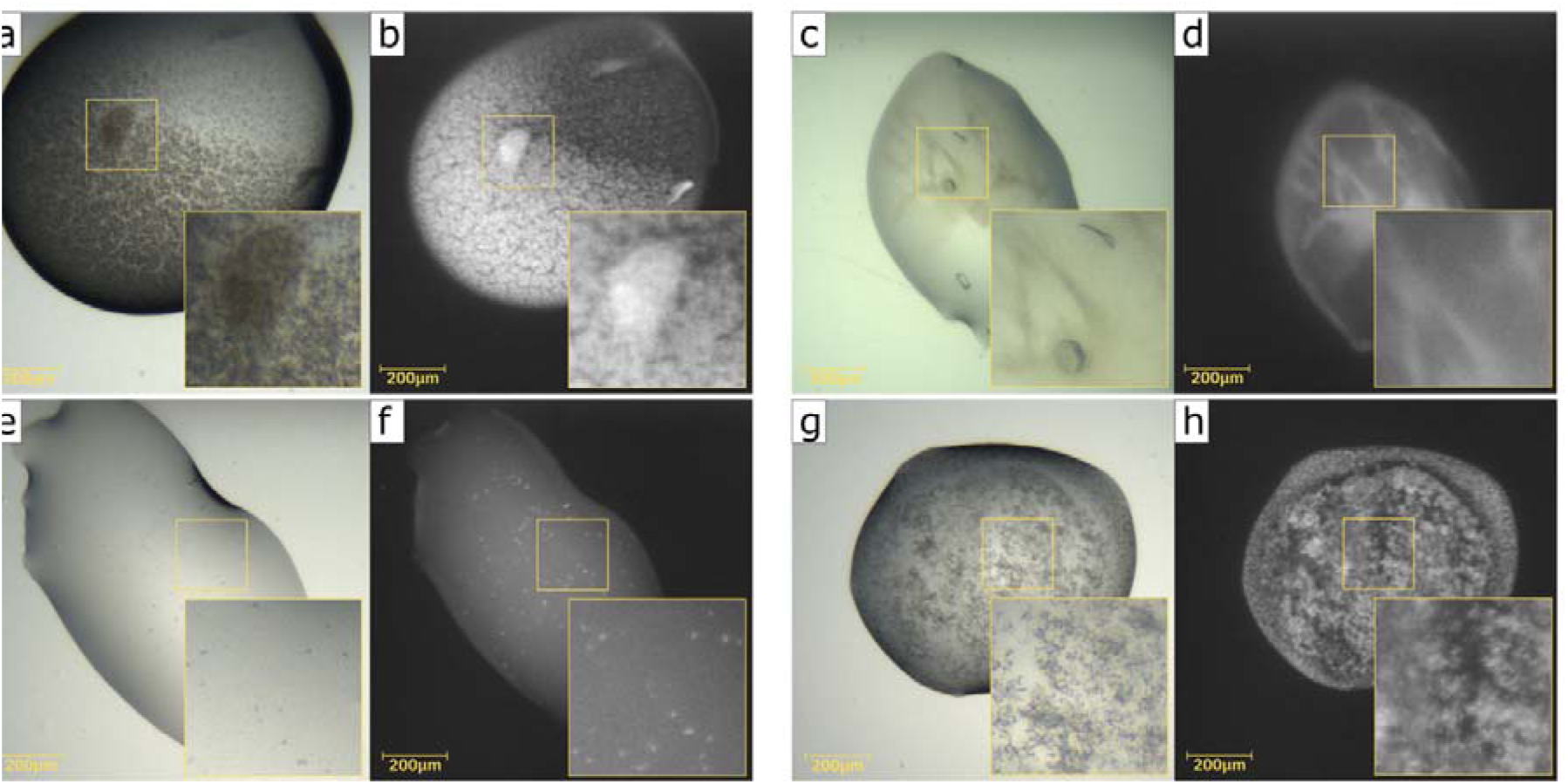
Comparison of the outcomes of Visible and UV light classifiers. Examples of crystals leading to different outcomes when evaluated by the Visible and UV model classifiers are shown. a-b, c-d, e-f and g-f panels show the same crystallization experiments images under visible (a-c, e-g) and UV light (b-d, f-h). Crystals in the a-b, e-f image pairs were only identified by the UV classifier. Crystals in the c-d, g-h image pairs were identified only by the Visible light classifier.

These results suggested that although showing lower overall performance, the AXIS-UV-v1 model can identify crystals under conditions where the visible model would fail and *vice versa*. Therefore, we set out to explore whether the AXIS-Vis-v1 and AXIS-UV-v1 models could be combined to provide optimal performance. We studied different ways of combining the numerical output of both models to produce a single consolidated score that would represent the likelihood of the experiment to contain crystals, including the arithmetic mean, product and *arguments of the Maxima*, or Argmax, which in this case consists in using as final score the outcome from the classification model that provides the highest crystal probability. As can be appreciated in Table 3, the approach that performed better was the Argmax. By applying the Argmax transformation we obtained the model AXIS-CRIMS-v1 classifier. In this model, both visible and UV images from the same crystallization experiment are independently evaluated and the numerical outcome from the model showing the highest crystal probability is retained as the final score. If no UV images are available, as is the case for the (C3 test dataset) the outcome of the visible classifier is directly used. Table 2 shows that the AXIS-CRIMS-v1 classifier improved performance as compared to the classifier based on either visible or UV images only (Table 3) with a notable increase in crystal recall, 95.54% and a balanced accuracy of 91.87% on the CRIMS test set. This demonstrates that the inclusion of UV images can improve significantly the performance of crystal image classification helping recall crystals that are difficult to identify with visible images only.

### 3.3. Continuous learning through a Lab-In-The-Loop approach

As shown above and previously reported (Rosa *et al*., 2023) inclusion of local images can have a strong impact in the accuracy of ML models for the classification of crystallization images. We reasoned that continuous addition of local images to the training process and particularly those where the classifier models have failed, could help improve performance. However, many laboratories do not have the capacity or resources to prepare and annotate extensive datasets from local image collections needed to re-train the models, which has limited the general use of crystal classification tools. We took advantage of the expert-driven CRIMS software (Cornaciu *et al*., 2021) to implement a Lab-In-The-Loop approach into AXIS enabling continuous interaction between human and AI annotations for continued ML training.

CRIMS is a web-based software suite that provides interfaces for experimental design as well as automated data and metadata tracking over the entire protein-to structure workflow (Cornaciu *et al*., 2021). CRIMS is built as an interactive tool that provides access to experimental results in real time and can collect input from hundreds of expert users (Cornaciu *et al*., 2021; Healey *et al*., 2021). It provides web-based interfaces for the screening and optimisation of crystallization experiments, communicates with crystallization and crystal imaging robots and automatically presents crystallization images to users for evaluation. Additional CRIMS modules enable automated crystal harvesting and communication with synchrotrons for automated diffraction data collection enabling seamless data and metadata exchange as well as recovery of diffraction data for downstream data processing (Cornaciu *et al*., 2021; Münzker *et al*., 2020; Healey *et al*., 2021; Zander *et al*., 2016). A dedicated CRIMS interface connects to the output of the automated crystal farms, presents crystallization images to users and enables them to record their own scores over a whole crystallization plate and over multiple plates (Cornaciu *et al*., 2021). We decided to exploit CRIMS to capture input from expert crystallographers distributed across multiple countries using the online crystallography services provided by HTX Lab in Grenoble and compare it to the output of automated annotations provided by AXIS. For this automated image scoring with the AXIS-CRIMS-v1 model was integrated into the CRIMS workflow for all new images produced by our crystal farms. The system takes about 150ms to infer the presence of crystals on a pair of visible and UV light images, making it possible to annotate 500 000 experiments per day on a dedicated GPU (see materials and methods section). The AXIS scores were recorded into the CRIMS databases and presented to scientists through the main CRIMS crystallization plate interface for expert evaluation. This interface presents images of each crystallization experiment in a microplate plate along with a map of the microplate to facilitate navigation from one experiment to the next. AXIS scores are inserted in the plate map and shown as different shades of green. To avoid overwhelming users with low probability predictions, we used a non-linear colour scale, giving very low visibility to probabilities below 40% and a colour intensity growing rapidly beyond (Figure 3). Moving the mouse over a specific experiment shows the crystal probability as a numerical value. While the AXIS scores are shown in this interface by default, users can introduce their own manual scores, which are stored independently by CRIMS and presented as a different colour code. This allowed us to collect feedback from expert crystallographers in a convenient way and compare it with the output of AXIS. Figure 3 shows plate maps corresponding to representative crystallization plates with AXIS scores and manual scores from crystallographers. As can be observed there is good agreement between them. However, there are also differences. Careful inspection of the discrepancies showed that they were in part associated with the level of performance of the ML classifier but in other cases they were caused by inaccuracies in the scores from human experts. As discussed above, inconsistencies of human scores are a well-known issue. This made it necessary to introduce a curation process before this data could be used for re-training.

**Figure 3.**
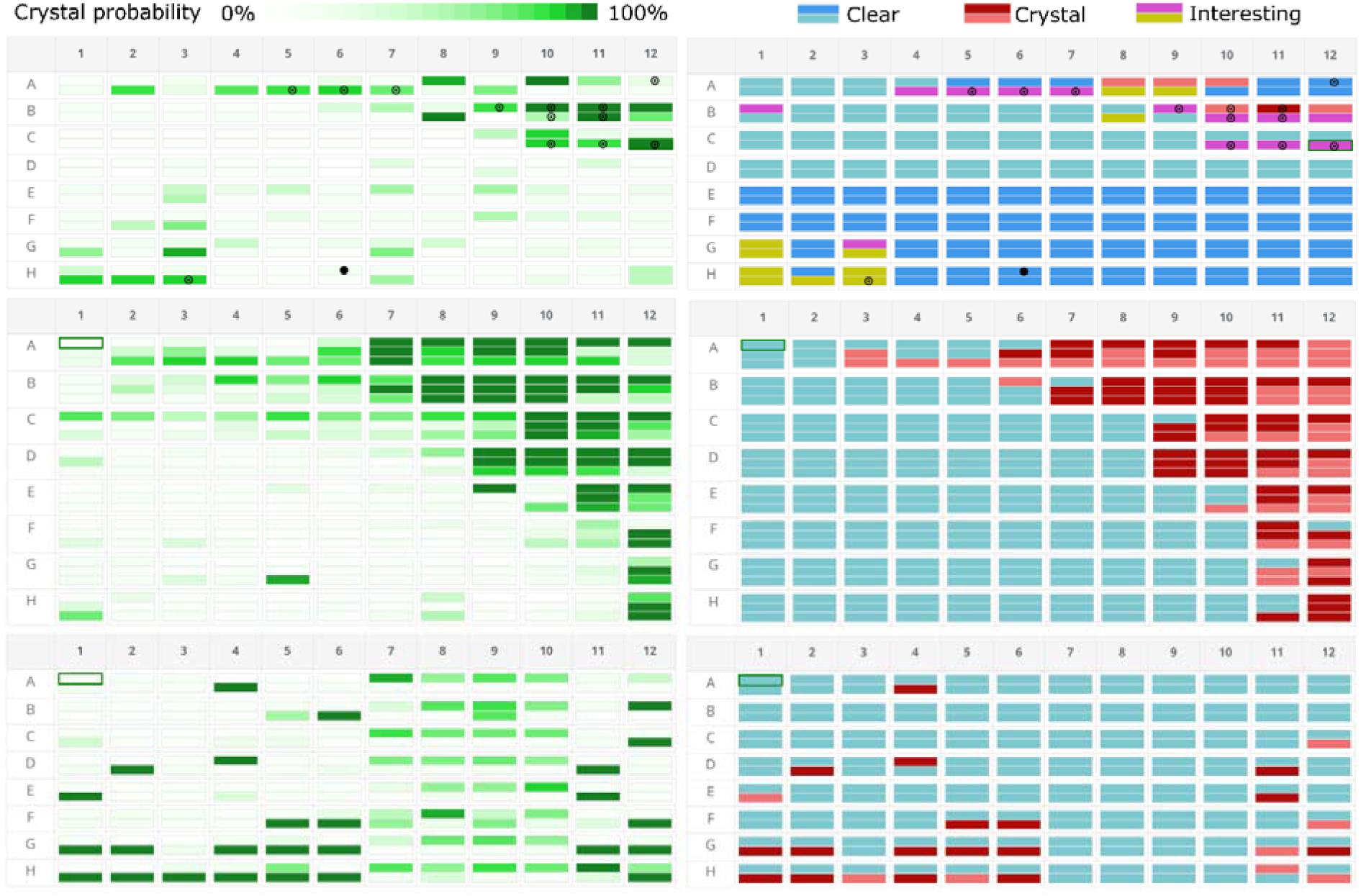
Comparison of ML and expert annotations. Dedicated CRIMS interfaces provide web-access for expert crystallographers to the outcomes of the AXIS image scores along with the crystallization images (not shown here). Crystal probabilities, as calculated by AXIS, are represented in different shades of green over a full 96-well crystallization plate (Left, panels corresponding to three different plates are shown). At the same time crystallographers can introduce their own expert scores online presented as a colour code from blue to dark red (Right). Good agreement between AXIS and expert inputs can be appreciated indicating the performance of the AXIS software, but few discrepancies can also be observed, for example position H11-3 in the middle panel.

In order to implement a Lab-In-The-Loop approach and evaluate discrepancies between AXIS and human scorers we built an AXIS dataset curation tool in CRIMS. The goal was to facilitate collection and annotation of datasets at any given facility. This tool automatically identifies discrepancies between human and AXIS scores and presents them through a web interface for manual evaluation by one or several expert curators (Figure 4). Both visible and UV images (if available) are presented in this interface along with the crystal probability provided by AXIS and the manual score. The CRIMS curator can apply one of 4 labels to the images: *Crystal*, *Other*, *Uncertain* (if the curators is uncertain) and *Unusable* (if the image shows any technical defect that prevents evaluation). Images marked as *Crystal* or *Other* by curators are then collected along with those for which the human and AXIS scores agreed to form a new training dataset.

**Figure 4.**
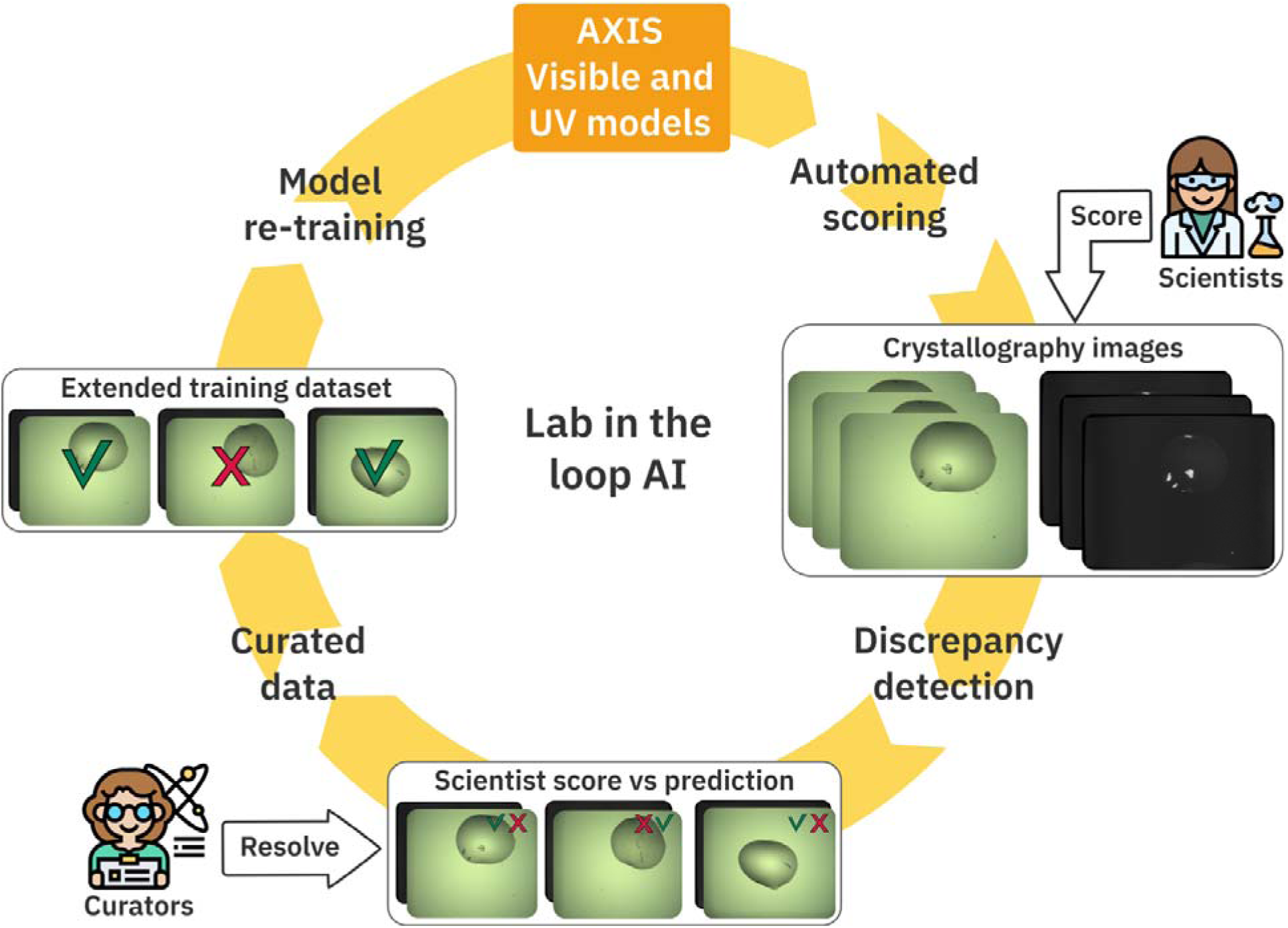
A Lab-In-The-Loop approach combining AI and Expert Input for continuous Machine Learning. The operation of recursive Lab-In-The-Loop cycles for continuous ML is presented. The AXIS foundation model is applied by CRIMS to assign crystal probabilities to all crystallization images produced at the local facility. AXIS scores, along with the images are present to expert crystallographers using the facility, via CRIMS web interfaces, that can introduce their own annotations. Discrepancies between ML and human scores are automatically collected, curated and assembled into a new training dataset to produce improved AI models. This cycle can be operated as many times as necessary to achieve optimal performance and continuous adaptation to the changing conditions at any particular facility.

A first set of 14.400 images were subjected to the AXIS Lab-In-the-Loop workflow. This generated a total of 1.140 experiments with discrepancies between human and AXIS scores. About half of the discrepancies (552) were experiments that the model wrongly identified as crystals. However, there were 17 cases where AXIS correctly identified crystals that the user had clearly missed. An additional 68 experiments contained crystals missed by AXIS but correctly identified by the users. The rest of the discrepancies (428) corresponded to unsolvable conflicts, either through lack of agreement among curators or due to image quality issues and were excluded from the curated dataset. This workflow resulted in the curated CRIMS-v2 training dataset consisting of 25.568 visible images and 16.658 UV images (Table 1).

The curated dataset described above was used to re-train the AXIS-Vis-v1 and AXIS-UV-v1models as described above and the output of both modes was again combined through Argmax to produce the AXIS-CRIMS-v2 classifier model (Figure 4). This new model was tested against the C3 and CRIMS training datasets. The performance of the classifier after the Lab-In-The-Loop training cycle improved notably both with the local and C3 test datasets, as show in Table 2. Supplementary Figure S1 shows confusion matrices for the different classifier models indicating the major area of improvement after this first Lab-In-The-Loop step was in precision, with 367 drops wrongly classified as crystals before this step and only 165 after it. This is not surprising, since crystallization datasets are largely imbalanced with a majority of images corresponding to the *other* class (73% of the images in the case of the AXIS-CRIMS-v2 training dataset). The AXIS-CRIMS-v2 classifier model was put in production to classify fresh experiments as they were been produced at the HTX lab and a few weeks later a new Lab-In-The-Loop training cycle took place (Figure 4). This time the new curated training set (CRIMS-v3, in table 1) contained 167.990 images and was used to re-train the AXIS-CRIMS-v2 model producing the new classifier model AXIS-CRIMS-v3. As indicated in table 2, the AXIS-CRIMS-v3 has a crystal recall of 96.43 % with the CRIMS-test dataset and a balanced accuracy of 96.27 % and is the one currently in production at the HTX lab.

The performance of AXIS classifiers is not homogeneous across all crystal types (Table 5). It shows very high detection rates for both small and medium size crystals, needles and clusters (above 98% recall) and excellent results with single crystals, while it shows lower performance with micro-crystals (92 %). In fact, 7 out of 8 false negatives produced by the AXIS-CRIMS-v3 model, correspond to very fine microcrystalline precipitates. The microcrystalline nature of this type of material is sometimes difficult to judge from a single image and often expert crystallographers would disagree as to whether such material should be classified as crystalline. Therefore, this class tends to be underrepresented in crystallization image test sets. However, we decided to include this category as crystalline and train AXIS to identify this type of material, because these experiments can sometimes provide useful information for follow-up crystal optimisation or be useful for other techniques, like serial crystallography or electron diffraction for example. As can be observed from Table 2. The AXIS-CRIMS v1, v2 and v3 models perform better that the MARCO model with all test sets. However, they tend to show slightly better crystal recall with the C3-test dataset than with the CRIMS-test set. This can be explained by the higher abundance of microcrystals in the CRIMS test dataset as compared to the MARCO and C3 test sets.

**Table 5.** AXIS-CRIMS recall scores per crystal types on CRIMS-test set.

| Model | Micro-crystals | Needles & Clusters | Single crystals |
| --- | --- | --- | --- |
| MARCO | 57.14% | 85.5% | 58.06% |
| AXIS-foundation | 74.49% | 89.47% | 90.32% |
| AXIS-CRIMS-v1 | 92.30% | 98.68% | 100.00% |
| AXIS-CRIMS-v2 | 92.86% | 98.68% | 100.00% |
| AXIS-CRIMS-v3 | 93.88% | 98.68% | 100.00% |

### 3.4. Extending AXIS to other image datasets

In order to determine whether AXIS could be transferable to image datasets generated at other facilities, we applied the same approach to a dataset generated at the VMXi beamline of the Diamond Light Source (Sanchez-Weatherby *et al*., 2019a; Mikolajek *et al*., 2023). The VMXi dataset contains 11 161 images in the same four classes defined for the MARCO dataset and an accompanying test set with 632 images (King *et al*., 2024). The images were obtained with a Rock Imager instrument (Formulatrix, Bedford, MA, United States) installed at the VMXi beamline using only visible light and different crystallization plate types and imaging conditions (King *et al*., 2024) as compared to those used at the HTX Lab facility in Grenoble. For example, in the VMXi dataset not only the crystallization drop, but also the whole crystallization well is visible in the micrographs. The VMXi training and test datasets were consolidated into two classes: “crystals”, “other” used by AXIS as indicated above.

We re-trained the AXIS-Foundation model with the VMXi training set using the same protocols described above in the materials and methods section. This produced the AXIS-VMXi classifier. We evaluated the performance of AXIS-VMXi against the VMXi test datasets. For comparison the MARCO (Bruno *et al*., 2018) and CHIMP v2 (King *et al*., 2024) classifier models, previously applied to VMXi data, were also run against the test dataset. As shown in Table 6, The AXIS-VMXi model produced very good performance against the VMXi test set, with crystal recall of 94.48% and balanced accuracy of 92.01%, improving both over the MARCO and CHIMPv2 models. This demonstrates that AXIS can be efficiently applied to the classification of crystallization images from different origins and under varying conditions with high performance.

**Table 6.**
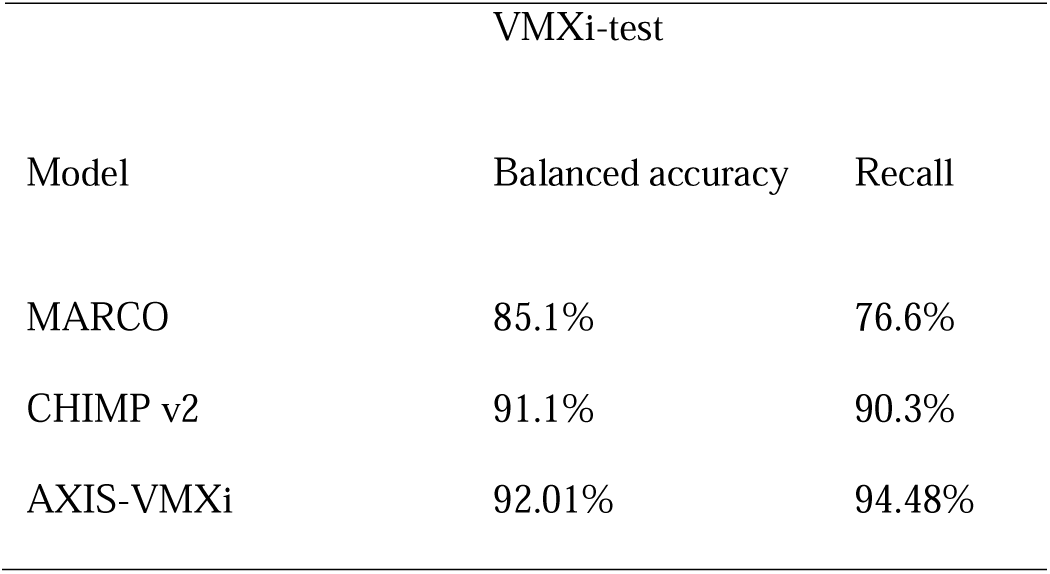
Extending AXIS to the VMXi dataset.

| VMXi-test |  |  |
| --- | --- | --- |
| Model | Balanced accuracy | Recall |
| MARCO | 85.1% | 76.6% |
| CHIMP v2 | 91.1% | 90.3% |
| AXIS-VMXi | 92.01% | 94.48% |

**Table 7.** Training complexity comparison.

| Model | Architecture | Total parameters | Trained parameters | Image inferences in training |
| --- | --- | --- | --- | --- |
| MARCO | Inception v3 | 24M | 24M | 100M |
| C3 | ResNet 50 | 25M | 25M | 22M |
| CHIMP v2 | ConvNext tiny | 28.5M | 28.5M | 5M |
| Axis-foundation | DINOv2-base | 86.6M | 4M | 900k |
| AXIS-CRIMS-v1 | DINOv2-base | 86.6M | 4M | 16k |
| AXIS-CRIMS-v2 | DINOv2-base | 86.6M | 4M | 50k |
| AXIS-CRIMS-v3 | DINOv2-base | 86.6M | 4M | 600k |

## 4. Discussion

Since the introduction of high throughput crystallization, automated crystal identification has been pursued (Rosa *et al*., 2023; Wilson, 2002; Bern *et al*., 2004; Saitoh *et al*., 2005) proving to be a difficult task. With the recent generalisation of automated protein-to-structure and ligand screening pipelines, capable of processing hundreds to thousands of crystals within a few days, reliable systems for automated crystal identification able to work with images from diverse origins become a necessity. Such systems have the potential of removing manual steps in otherwise highly automated pipelines increasing their productivity and reliability with impact both in fundamental research and structure-based drug design. ML-based classifiers for crystallization images have been developed in a number of laboratories (Bruno *et al*., 2018; King *et al*., 2024), but their performance tend to degrade when applied to images from other facilities (Rosa *et al*., 2023). This imposes the need of re-training with image datasets produced locally, thereby limiting their transferability. The AI-based Crystal Identification System (AXIS) described here, addresses these limitations by integrating recent foundational models for computer vision, state-of-the-art transfer learning techniques and a web-based Lab-In-the-Loop approach combining AI and collaborative expert input for continuous learning, to deliver high performance in crystal detection and enable rapid adaptation to local or varying conditions at any facility.

AXIS applies a multi-step ML approach (Figure 1) and provides a foundational model for crystal identification able to interpret both visible and UV light images. The AXIS-Foundation model integrates the DINOv2-base computer vision model (Oquab *et al*., 2023) trained to extract visual features using 142 million images and has been re-trained for crystal identification using the MARCO (Bruno *et al*., 2018) dataset. This represents an ideal starting model for crystal identification as it combines the power of a large computer vision model with the largest crystallization image dataset available to date, with 493,214 crystallization images originating from 5 different laboratories (Bruno *et al*., 2018). The AXIS-foundation model performs better than previously reported models across different test image datasets, therefore it is an ideal starting point for crystal image classification. It can be easily implemented at any facility and requires minimal computing resources as it is able to process hundreds of thousands of images per day in a single GPU. At the same time, AXIS performance at any given facility is likely to improve by fine-tuning with image datasets generated locally. Collection and annotation of image datasets can be a tedious and time-consuming process that many laboratories might find difficult to implement. By integrating AXIS with the CRIMS software (Cornaciu *et al*., 2021) we have implemented a Lab-In-The-Loop approach that facilitates this operation significantly lowering the barriers for the implementation of AI in crystal detection.

The CRIMS Lab-In-the-Loop functionality automatically tracks ML-based and human image annotations during the normal progress of crystallography projects and collects agreements and discrepancies. A data curation interface enables efficient evaluation of the data by expert crystallographers to eliminate human annotation inconsistencies and generate validated image datasets that can be used for fine-tuning of the initial model under local conditions. This approach makes it possible to integrate input from hundreds of crystallographers over a very diverse range of samples making the collection of local image datasets very efficient and saving expert’s worktime. The fine-tuning process can be applied in multiple-steps and with limited size datasets, which facilities implementation. For example, the AXIS-foundational model can be implemented as a first step in crystal detection while at the same time it will provide the basis for comparison between user input and ML-based scores under local conditions. After a few weeks, sufficient data would be available for a first fine-tuning step that can then be repeated until optimal performance is achieved. As we demonstrate here this approach was successful when applied to images from two different facilities, the HTX lab at EMBL Grenoble (Cornaciu *et al*., 2021) and the Diamond VMXi beam line (Sanchez-Weatherby *et al*., 2019a), showing that combination of AXIS-foundation with Lab-In-The-Loop approach represents an efficient way to achieved high level of crystal detection as well as accuracy with datasets from different origins. At the HTX facility where the system is currently in production, the AXIS-CRIMS-v3 model trained through consecutive Lab-in-the-loop cycles, achieved very high performance in the identification of 2-d and 3-d crystals (crystal needles, plates, clusters, and single crystals) with over 98 % detection and can also identify difficult categories, like microcrystalline precipitates with a recall of 94%.

We also demonstrate that the use of UV images can improve crystal detection. Currently, automated crystallization plate imagers are often equipped with both visible and UV imaging capabilities, but UV imaging in AI-based crystal detection had not been systematically explored to date. Our results show that when addressed independently, the performance of the UV-based image classifier is lower compared to the visible image model. This may be in part due to the fact that the UV classifier has been trained with a comparatively lower number of images, as large and diverse UV-image datasets are still lacking. However, the UV-based classifier was able to identify crystals that the visible model failed to recognise. Therefore, the combination of both visible and UV image models improves the performance of image classification. In its current implementation, AXIS is able to use either visible or UV images to provide scores. However, if both type of images are available for the same experiment it will automatically combine the scores to optimal results. Thanks to this approach AXIS achieves a performance comparable to that of expert crystallographers but in a fully automated manner. Moreover, it eliminates many of the problems associated with manual annotation, particularly inconsistencies due to fatigue or lack of experience for example. Indeed, AXIS regularly identifies crystals that have been missed by human inspection, helping scientists identify key crystallization conditions while at the same time their expert input reinforces the machine learning process.

Potential applications of AXIS extend beyond initial crystal identification and could contribute to automation in other areas of the crystallography workflow. For example, automated crystal centring through X-ray sample raftering is currently used at many synchrotron beam lines. However, it consumes time potentially slowing down the data collection process. Systems like AXIS have the potential to replace sample rastering making data collection much more efficient and cost-effective at synchrotrons. On the other hand, the combination of ML and Lab-In-The-Loop approach we demonstrate here can be applied to other areas of structural biology were initial ML-trained models can learn through input from expert scientists to choose optimal experimental parameters throughput very complex experimental workflows that would otherwise require careful expert evaluation and human decision. The approach we demonstrate here, in combination with existing laboratory automation can help transform a once considered complex and time consuming experimental workflow available only to well-trained experts into fully automated workflows where complex experimental parameters are automatically chosen to achieve results with optimal quality making structural biology facilities worldwide more efficient and helping shift scientists time from experiment control to data analysis and interpretation.

## Synopsis

AXIS, **A**I-based **C**rystal **I**dentification **S**ystem combines artificial intelligence with an iterative Lab-in-the-loop approach for automated crystal detection. AXIS ensures high performance in crystal detection, requires small size training datasets and facilitates rapid adaptation to local and changing conditions at any given facility.

## Acknowledgements

We are grateful to Anna Kreschuk (EMBL) for advice and support, Peter Murphy and Fynn Beuttenmueller from EMBL for initial exploratory work, Jan Korbel and the EMBL Data Science Centre as well as Rupert Lück EMBL IT Team for support and access to HPC resources. We want to acknowledge access to the EMBL-ESRF Joint Structural Biology and Imaging Beamlines at ESRF. This project received financial support from the European Commission through EU project Fragment-Screen, grant agreement ID: 101094131. AP received an EMBL ARISE fellowship supported by the European Union’s Horizon 2020 research and innovation programme under the Marie Skłodowska-Curie grant agreement No 945405.

## Data availability

Training and inference scripts are available at https://github.com/marquez-group-embl/AXIS. Training and test datasets are available from zenodo.org, See table 1. Trained models are available at https://huggingface.co/Marquez-Group-EMBL.

## Appendix B. Supporting information

**S1. Supplementary Figure 1:**
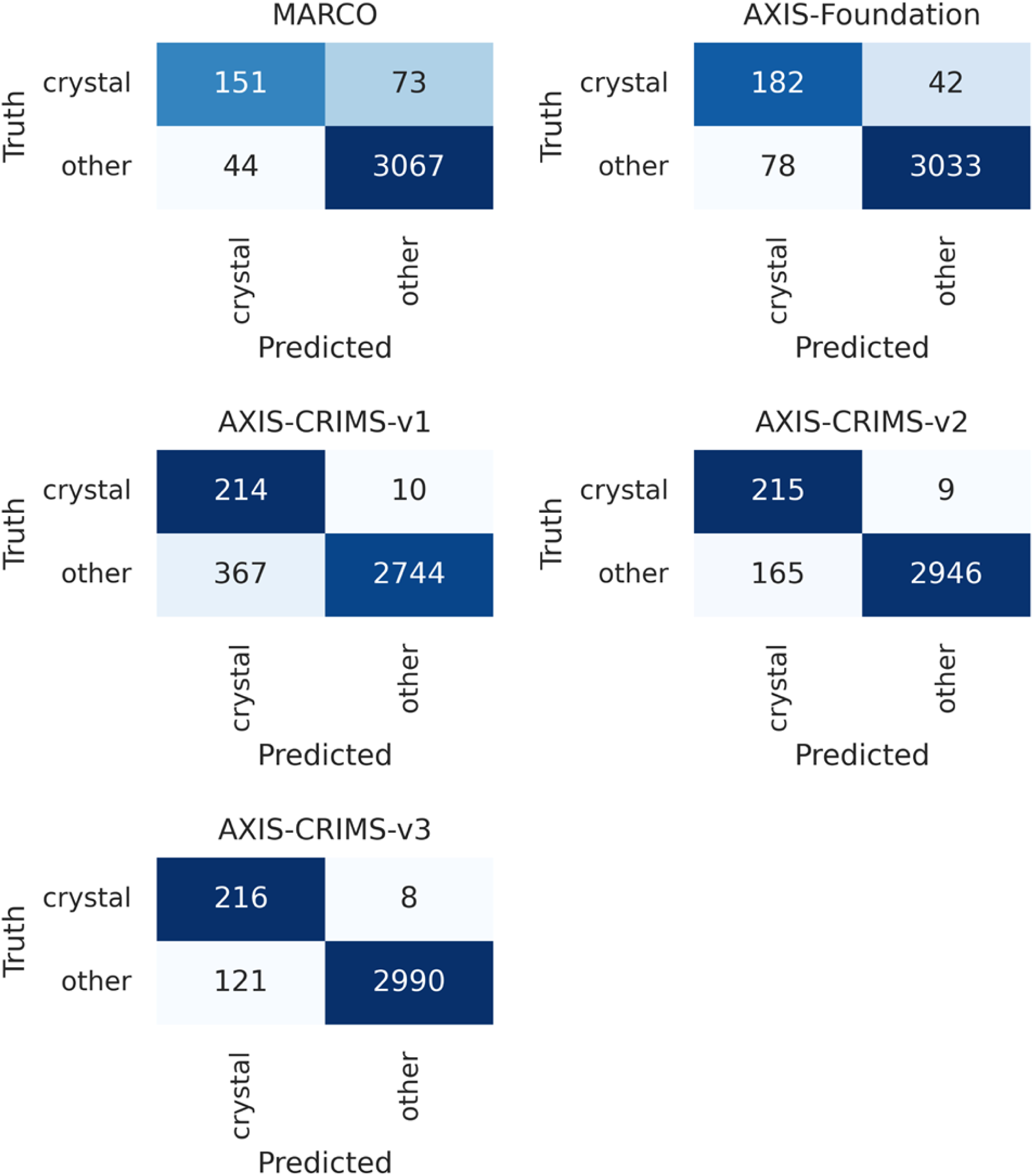
Confusion matrices of the different AXIS classifiers with the CRIMS-test dataset

## References

Abramson, J., Adler, J., Dunger, J., Evans, R., Green, T., Pritzel, A., Ronneberger, O., Willmore, L., Ballard, A. J., Bambrick, J., Bodenstein, S. W., Evans, D. A., Hung, C.-C., O’Neill, M., Reiman, D., Tunyasuvunakool, K., Wu, Z., Žemgulytė, A., Arvaniti, E., Beattie, C., Bertolli, O., Bridgland, A., Cherepanov, A., Congreve, M., Cowen-Rivers, A. I., Cowie, A., Figurnov, M., Fuchs, F. B., Gladman, H., Jain, R., Khan, Y. A., Low, C. M. R., Perlin, K., Potapenko, A., Savy, P., Singh, S., Stecula, A., Thillaisundaram, A., Tong, C., Yakneen, S., Zhong, E. D., Zielinski, M., Žídek, A., Bapst, V., Kohli, P., Jaderberg, M., Hassabis, D. & Jumper, J. M. (2024). Nature 630, 493–500.

Baek, M., DiMaio, F., Anishchenko, I., Dauparas, J., Ovchinnikov, S., Lee, G. R., Wang, J., Cong, Q., Kinch, L. N., Schaeffer, R. D., Millán, C., Park, H., Adams, C., Glassman, C. R., DeGiovanni, A., Pereira, J. H., Rodrigues, A. V., Van Dijk, A. A., Ebrecht, A. C., Opperman, D. J., Sagmeister, T., Buhlheller, C., Pavkov-Keller, T., Rathinaswamy, M. K., Dalwadi, U., Yip, C. K., Burke, J. E., Garcia, K. C., Grishin, N. V., Adams, P. D., Read, R. J. & Baker, D. (2021). Science 373, 871–876.

Barthel, T., Benz, L., Basler, Y., Crosskey, T., Dillmann, A., Förster, R., Fröling, P., Dieguez, C. G., Gless, C., Hauß, T., Hellmig, M., Jänisch, L., James, D., Lennartz, F., Mijatovic, J., Oelker, M., Scanlan, J. W., Weber, G., Wollenhaupt, J., Mueller, U., Dobbek, H., Wahl, M. C. & Weiss, M. S. (2024). Applied Research 3, e202400110.

Bern, M., Goldberg, D., Stevens, R. C. & Kuhn, P. (2004). J Appl Crystallogr 37, 279–287.

Bowler, M. W., Nurizzo, D., Barrett, R., Beteva, A., Bodin, M., Caserotto, H., Delagenière, S., Dobias, F., Flot, D., Giraud, T., Guichard, N., Guijarro, M., Lentini, M., Leonard, G. A., McSweeney, S., Oskarsson, M., Schmidt, W., Snigirev, A., von Stetten, D., Surr, J., Svensson, O., Theveneau, P. & Mueller-Dieckmann, C. (2015). J Synchrotron Rad 22, 1540– 1547.

Bruno, A. E., Charbonneau, P., Newman, J., Snell, E. H., So, D. R., Vanhoucke, V., Watkins, C. J., Williams, S. & Wilson, J. (2018). PLoS ONE 13, e0198883.

Chayen, N. E. & Saridakis, E. (2008). Nat Methods 5, 147–153.

Cipriani, F., Felisaz, F., Launer, L., Aksoy, J.-S., Caserotto, H., Cusack, S., Dallery, M., di-Chiaro, F., Guijarro, M., Huet, J., Larsen, S., Lentini, M., McCarthy, J., McSweeney, S., Ravelli, R., Renier, M., Taffut, C., Thompson, A., Leonard, G. A. & Walsh, M. A. (2006). Acta Crystallogr D Biol Crystallogr 62, 1251–1259.

Cornaciu, I., Bourgeas, R., Hoffmann, G., Dupeux, F., Humm, A.-S., Mariaule, V., Pica, A., Clavel, D., Seroul, G., Murphy, P. & Márquez, J. A. (2021). *JoVE* 62491.

Cox, O. B., Krojer, T., Collins, P., Monteiro, O., Talon, R., Bradley, A., Fedorov, O., Amin, J., Marsden, B. D., Spencer, J., von Delft, F. & Brennan, P. E. (2016). Chem. Sci. 7, 2322–2330.

Cumbaa, C. A. & Jurisica, I. (2010). J Struct Funct Genomics 11, 61–69.

Cumbaa, C. A., Lauricella, A., Fehrman, N., Veatch, C., Collins, R., Luft, J., DeTitta, G. & Jurisica, I. (2003). Acta Crystallogr D Biol Crystallogr 59, 1619–1627.

Cusack, S., Belrhali, H., Bram, A., Burghammer, M., Perrakis, A. & Riekel, C. (1998). Nat Struct Mol Biol 5, 634–637.

Devlin, J., Chang, M.-W., Lee, K. & Toutanova, K. (2018). 10.48550/ARXIV.1810.04805.

Dimasi, N., Flot, D., Dupeux, F. & Márquez, J. A. (2007). Acta Crystallogr F Struct Biol Cryst Commun 63, 204–208.

Dosovitskiy, A., Beyer, L., Kolesnikov, A., Weissenborn, D., Zhai, X., Unterthiner, T., Dehghani, M., Minderer, M., Heigold, G., Gelly, S., Uszkoreit, J. & Houlsby, N. (2020). 10.48550/ARXIV.2010.11929.

Healey, R. D., Basu, S., Humm, A.-S., Leyrat, C., Cong, X., Golebiowski, J., Dupeux, F., Pica, A., Granier, S. & Márquez, J. A. (2021). An automated platform for structural analysis of membrane proteins through serial crystallography Biochemistry.

Helliwell, J. R. (2017). Bioscience Reports 37, BSR20170204.

Hu, E. J., Shen, Y., Wallis, P., Allen-Zhu, Z., Li, Y., Wang, S., Wang, L. & Chen, W. (2021a). 10.48550/ARXIV.2106.09685.

Hu, E. J., Shen, Y., Wallis, P., Allen-Zhu, Z., Li, Y., Wang, S., Wang, L. & Chen, W. (2021b). 10.48550/ARXIV.2106.09685.

Jumper, J., Evans, R., Pritzel, A., Green, T., Figurnov, M., Ronneberger, O., Tunyasuvunakool, K., Bates, R., Žídek, A., Potapenko, A., Bridgland, A., Meyer, C., Kohl, S. A. A., Ballard, A. J., Cowie, A., Romera-Paredes, B., Nikolov, S., Jain, R., Adler, J., Back, T., Petersen, S., Reiman, D., Clancy, E., Zielinski, M., Steinegger, M., Pacholska, M., Berghammer, T., Bodenstein, S., Silver, D., Vinyals, O., Senior, A. W., Kavukcuoglu, K., Kohli, P. & Hassabis, D. (2021). Nature 596, 583–589.

King, O. N. F., Levik, K. E., Sandy, J. & Basham, M. (2024). Acta Crystallogr D Struct Biol 80, 744– 764.

Krishna, R., Wang, J., Ahern, W., Sturmfels, P., Venkatesh, P., Kalvet, I., Lee, G. R., Morey-Burrows, F. S., Anishchenko, I., Humphreys, I. R., McHugh, R., Vafeados, D., Li, X., Sutherland, G. A., Hitchcock, A., Hunter, C. N., Kang, A., Brackenbrough, E., Bera, A. K., Baek, M., DiMaio, F. & Baker, D. (2024). Science 384, eadl2528.

Liu, R., Freund, Y. & Spraggon, G. (2008). Acta Crystallogr D Biol Crystallogr 64, 1187–1195.

Liu, Z., Mao, H., Wu, C.-Y., Feichtenhofer, C., Darrell, T. & Xie, S. (2022). 10.48550/ARXIV.2201.03545.

McCarthy, A. A., Barrett, R., Beteva, A., Caserotto, H., Dobias, F., Felisaz, F., Giraud, T., Guijarro, M., Janocha, R., Khadrouche, A., Lentini, M., Leonard, G. A., Lopez Marrero, M., Malbet-Monaco, S., McSweeney, S., Nurizzo, D., Papp, G., Rossi, C., Sinoir, J., Sorez, C., Surr, J., Svensson, O., Zander, U., Cipriani, F., Theveneau, P. & Mueller-Dieckmann, C. (2018). J Synchrotron Rad 25, 1249–1260.

McPherson, A. (2017). Vol. 1607, Protein Crystallography, edited by A. Wlodawer, Z. Dauter & M. Jaskolski. pp. 17–50. New York, NY: Springer New York.

Mehrabi, P., Schulz, E. C., Agthe, M., Horrell, S., Bourenkov, G., Von Stetten, D., Leimkohl, J.-P., Schikora, H., Schneider, T. R., Pearson, A. R., Tellkamp, F. & Miller, R. J. D. (2019). Nat Methods 16, 979–982.

Mikolajek, H., Sanchez-Weatherby, J., Sandy, J., Gildea, R. J., Campeotto, I., Cheruvara, H., Clarke, J. D., Foster, T., Fujii, S., Paulsen, I. T., Shah, B. S. & Hough, M. A. (2023). IUCrJ 10, 420– 429.

Münzker, L., Petrick, J. K., Schleberger, C., Clavel, D., Cornaciu, I., Wilcken, R., Márquez, J. A., Klebe, G., Marzinzik, A. & Jahnke, W. (2020). ChemBioChem 21, 3096–3111.

Newman, J., Egan, D., Walter, T. S., Meged, R., Berry, I., Ben Jelloul, M., Sussman, J. L., Stuart, D. I. & Perrakis, A. (2005). Acta Crystallogr D Biol Crystallogr 61, 1426–1431.

Oquab, M., Darcet, T., Moutakanni, T., Vo, H., Szafraniec, M., Khalidov, V., Fernandez, P., Haziza, D., Massa, F., El-Nouby, A., Assran, M., Ballas, N., Galuba, W., Howes, R., Huang, P.-Y., Li, S.-W., Misra, I., Rabbat, M., Sharma, V., Synnaeve, G., Xu, H., Jegou, H., Mairal, J., Labatut, P., Joulin, A. & Bojanowski, P. (2023). 10.48550/ARXIV.2304.07193.

Pan, S., Shavit, G., Penas-Centeno, M., Xu, D.-H., Shapiro, L., Ladner, R., Riskin, E., Hol, W. & Meldrum, D. (2006). Acta Crystallogr D Biol Crystallogr 62, 271–279.

Rosa, N. & Newman, J. (2021). C3 Protein Crystallization Dataset Zenodo.

Rosa, N., Watkins, C. J. & Newman, J. (2023). PLoS ONE 18, e0283124.

Rupp, B., Segelke, B. W., Krupka, H. I., Lekin, T., Schäfer, J., Zemla, A., Toppani, D., Snell, G. & Earnest, T. (2002). Acta Crystallogr D Biol Crystallogr 58, 1514–1518.

Saitoh, K., Kawabata, K., Asama, H., Mishima, T., Sugahara, M. & Miyano, M. (2005). Acta Crystallogr D Biol Crystallogr 61, 873–880.

Sanchez-Weatherby, J., Sandy, J., Mikolajek, H., Lobley, C. M. C., Mazzorana, M., Kelly, J., Preece, G., Littlewood, R. & Sørensen, T. L.-M. (2019a). J Synchrotron Rad 26, 291–301.

Sanchez-Weatherby, J., Sandy, J., Mikolajek, H., Lobley, C. M. C., Mazzorana, M., Kelly, J., Preece, G., Littlewood, R. & Sørensen, T. L.-M. (2019b). Journal of Synchrotron Radiation 26, 291– 301.

Schlichting, I. (2015). IUCrJ 2, 246–255.

Schwalbe, H., Audergon, P., Haley, N., Amaro, C. A., Agirre, J., Baldus, M., Banci, L., Baumeister, W., Blackledge, M., Carazo, J. M., Carugo, K. D., Celie, P., Felli, I., Hart, D. J., Hauß, T., Lehtiö, L., Lindorff-Larsen, K., Márquez, J., Matagne, A., Pierattelli, R., Rosato, A., Sobott, F., Sreeramulu, S., Steyaert, J., Sussman, J. L., Trantirek, L., Weiss, M. S. & Wilmanns, M. (2024). Structure 32, 1563–1580.

Stauch, B. & Cherezov, V. (2018). Annu. Rev. Biophys. 47, 377–397.

Stock, D., Perisic, O. & Löwe, J. (2005). Progress in Biophysics and Molecular Biology 88, 311–327.

Thomas, S. E., Collins, P., James, R. H., Mendes, V., Charoensutthivarakul, S., Radoux, C., Abell, C., Coyne, A. G., Floto, R. A., Von Delft, F. & Blundell, T. L. (2019). Phil. Trans. R. Soc. A. 377, 20180422.

Vaswani, A., Shazeer, N., Parmar, N., Uszkoreit, J., Jones, L., Gomez, A. N., Kaiser, L. & Polosukhin, I. (2017). 10.48550/ARXIV.1706.03762.

Whittle, P. J. & Blundell, T. L. (1994). Annu. Rev. Biophys. Biomol. Struct. 23, 349–375.

Wilson, J. (2002). Acta Crystallogr D Biol Crystallogr 58, 1907–1914.

Yosinski, J., Clune, J., Bengio, Y. & Lipson, H. (2014). 10.48550/arXiv.1411.1792.

Yu, D., Chojnowski, G., Rosenthal, M. & Kosinski, J. (2023). Bioinformatics 39, btac749.

Zander, U., Hoffmann, G., Cornaciu, I., Marquette, J.-P., Papp, G., Landret, C., Seroul, G., Sinoir, J., Röwer, M., Felisaz, F., Rodriguez-Puente, S., Mariaule, V., Murphy, P., Mathieu, M., Cipriani, F. & Márquez, J. A. (2016). Acta Crystallogr D Struct Biol 72, 454–466.

Zhou, H.-Y., Lu, C., Yang, S. & Yu, Y. (2021). 10.48550/ARXIV.2108.05305.

Zhuang, F., Qi, Z., Duan, K., Xi, D., Zhu, Y., Zhu, H., Xiong, H. & He, Q. (2020). 10.48550/arXiv.1911.02685.

